# miR-199-3p Suppresses Cellular Migration and Viability and Promotes Progesterone Production in Goose Ovarian Follicles Before Selection by Targeting *ITGB8* and Regulating Other ECM-related Genes

**DOI:** 10.1101/2021.12.06.471452

**Authors:** Qin Li, Keshan Zhang, Xianzhi Zhao, Jing Li, Youhui Xie, Hang Zhong, Qigui Wang

**Affiliations:** Chongqing Academy of Animal Science, Rongchang county, Chongqing 402460, P. R. China; Chongqing Engineering Research Center of Goose Genetic Improvement, Chongqing, P. R. China

**Keywords:** *Anser cygnoides*, miR-199-3p, granulosa cells, cellular viability, cell migration, progesterone production, *ITGB8*, extracellular matrix

## Abstract

The extracellular matrix (ECM) constitutes the follicular basal lamina and is also present between follicular cells. Remodeling of the ECM is believed to be a key event in follicular development, especially follicular selection, and plays important roles in cell migration, survival, and steroidogenesis. miR-199-3p is differentially expressed in the goose follicular granulosa layer during follicular selection and is reported to play a primary role in inhibiting cell migration and invasion. Nevertheless, the effect of miR-199-3p on ovarian follicles and its role in follicular cellular migration are not understood. In this study, we demonstrated by qRT-PCR that miR-199-3p was differentially expressed in the granulosa layer from goose ovarian follicles before and after follicular selection. Additionally, we found that miR-199-3p overexpression could significantly suppress cell viability and migration, as well as elevate both the concentration of progesterone and the expression of key progesterone production genes in cultured granulosa cells (GCs) from goose pre-hierarchical follicles. Furthermore, using dual-fluorescence reporter experiments on 293T cells, we confirmed that miR-199-3p downregulated the expression of the ECM gene *ITGB8* by directly targeting its mRNA three prime untranslated region (3′ UTR). Finally, we found that miR-199-3p overexpression in the GCs of goose pre-hierarchical follicles inhibited the expression of two ECM-related genes (*MMP9* and *MMP15*) yet promoted the expression of another two ECM-related genes (*COL4A1* and *LAMA1*). Taken together, these findings suggest that miR-199-3p participates in granulosa cell migration, viability, and steroidogenesis in goose ovarian follicles before selection by targeting *ITGB8* and modulating other ECM-related genes. These data highlight the key roles of miR-199-3p in follicular cell migration, viability, and steroidogenesis by regulating ECM-related genes and thus contribute to a better understanding of the mechanisms underlying follicle selection in birds.

## Introduction

Most domestic goose breeds lay 30-40 eggs a year, which is far less than other poultry such as chickens and ducks (>300) [1]. The egg yield of poultry depends on the development and maturation of ovarian follicles, which is a complex process involving follicular recruitment, growth, selection, and dominance [2, 3]. Follicle selection, defined as the process by which single pre-hierarchical small yellow follicles (SWFs) are selected into hierarchical follicles, is widely accepted as the rate-limiting step of avian reproductive potential [4]. The dynamic selection of ovarian follicles is accompanied by cyclical remodeling and breakdown of the follicular wall, thus requiring extensive turnover of the extracellular matrix (ECM) [5, 6].

ECM, multiple proteins that constitute the follicular basal layer and exist also between follicular cells, can provide rigid or elastic mechanical support for tissues [7]. Collagens, laminins, fibronectin, and matrix metalloproteinases (MMPs) are the main components of ECM and take part in regulating granulosa cell (GC) behavior and function including migration, differentiation, survival, and steroidogenesis [8-12]. MMPs in particular play a crucial role in the breakdown of ECM substrates [10, 13, 14] and the regulation of cell migration in various tissue types [13, 15]. In the chicken, it has been suggested that *MMP2* and *MMP9* participate in the ECM remodeling that is required for follicular development [16]. Besides, laminins, fibronectin and laminin-integrin interactions enhance survival and proliferation and modulate estradiol (E2) secretion in ovine GCs [8, 17].

MicroRNAs (miRNAs) are non-coding RNA molecules with a length of 18-24 nucleotides. MiRNAs are involved in GC functions including cell proliferation, migration, steroidogenesis, and apoptosis by binding to the mRNA three prime untranslated region (3′ UTR) [18]. Our preliminary study profiling miRNA transcription in the granulosa layer of geese found that miR-199-3p is one of the microRNAs with the largest fold change (FC) during follicular selection [19]. Research by predecessors has shown that miR-199 can suppress cell migration, invasion, and proliferation by downregulating its target gene in various type of cancers, and is thus considered to function as a tumor suppressor and as a potential prognostic marker in cancer [20-22]. The miR-199 family, including miR-199a(b)-5p and miR-199a(b)-3p, inhibits cell migration and invasion in head, neck and bladder cancer by regulation of integrin α 3 (*ITGA3*) [20, 21]. miR-199a-3p reduces cell proliferation, G1 phase cell cycle arrest, cell invasion, and increases cell apoptosis in ovarian cancer by targeting integrin β8 (*ITGB8*) [23]. In short, miR-199 can regulate cell proliferation, apoptosis, and migration by targeting ECM-related genes. Although, miR-199 has been found to be expressed in the ovarian follicle of mammals and birds [19, 24, 25], the role of miR-199-3p in avian GCs, and whether miR-199-3p performs its role by targeting ITGBs or other ECM-related genes, is unknown.

In the present study, differential expression of miR-199-3p in the granulosa layer of pre-hierarchical follicles from geese during follicular selection was confirmed by qRT-PCR. Next, miR-199-3p was found to suppress viability and migration, as well as promote both the production of progesterone (P4) and key genes required for P4 production (*STAR* and *3βHSD*) in the cultured GCs from the pre-hierarchical follicles of geese. To understand the mechanism by which miR-199-3p operates, we identified the putative target gene of miR-199-3p, *ITGB8*, by combining a target gene prediction database with goose transcriptome data and further confirmed that miR-199-3p directly targets ECM-related gene *ITGB8* by luciferase reporter assay in 293T cells. In addition, we observed that miR-199a-3p overexpression resulted in a significant change in ECM-related gene expression (*MMP9, MMP15, COL4A1* and *LAMA1*) in cultured GCs from goose pre-hierarchical follicles. Taken together, these data demonstrate that miR-199-3p may be involved in viability, migration, and steroidogenesis of goose GCs by targeting *ITGB8* and regulating other ECM-related genes during avian follicular selection.

## MATERIALS AND METHODS

### Experimental animals and isolation of GCs

Healthy adult female Sichuan White geese (*Anser cygnoides*, 35–45 weeks old) from the Experimental Farm for Waterfowl Breeding at Chongqing Academy of Animal Science were used in the present study. Detailed husbandry information, laying cycle, and the classification criteria for ovarian follicles have been previously described [26]. Five geese were killed by cervical dislocation about 2 h before laying eggs. The granulosa layers of each individual were isolated from the pre-hierarchical (2-4, 4-6, 6-8 and 8-10 mm in diameter) and hierarchical (F5, F4, F3, F2, F1) follicles according to protocols described by Gilbert [27] and Deng [28]. Subsequently, the GCs isolated from each sized follicle (5 geese) were snap-frozen in liquid nitrogen and stored at -80°C until total RNA extraction. All experimental protocols involving animal manipulation were approved by the Laboratory Animal Management Committee of Chongqing Academy of Animal Sciences (Chongqing, China) under approval No. 2006-398.

### Culture and transfection of primary GCs and cell lines

GCs isolated from 6-10 mm diameter pre-hierarchical SWF follicles (5 geese) were used for primary cell culture and further *in vitro* cellular experiments. The procedure for primary GC culture was as follows: GCs of pre-hierarchical SWF follicles were digested by 0.25 % type II collagenase (Sigma-Aldrich, St. Louis, MO, USA) in a 37 °C water bath for approximately 3 min, and then incubated at 37°C under 5 % CO_2_ in Dulbecco modified eagle medium/Nutrient Mixture F-12 (1:1) (DMEM/F12; HyClone, Logan, UT, USA) containing 10 % fetal bovine serum (FBS) (Gibco, CA, USA) and a 1 % penicillin and streptomycin mixture (Gibco). Additionally, the human embryonic kidney 293T (HEK293T) cell line was obtained from the Kunming Institute of Zoology (Kunming, China) and incubated under the same conditions as primary GCs. MiR-199-3p mimic and mimic-Negative Control (NC) were double-stranded, whereas the inhibitor and inhibitor-NC were single-stranded. The sequences of miR-199-3p are shown in Supplementary Table S1 and were designed and synthesized by GenePharama (RIBOBio Co, Guangzhou, China). MiR-199-3p mimic and inhibitor were transfected into goose primary GCs or the HEK293T cell line for gain-of-function and loss-of-function experiments, respectively. The cellular experiments included cell viability and wound healing assays in goose primary GCs, as well as a dual luciferase reporter assay in HEK293T cells. Briefly, GCs or HEK293T cells were cultured in 12-well and 96-well plates. When the cells were approximately 75% confluent, small oligonucleotides were transfected at different concentrations using ViaFect^™^ Transfection Reagent (Promega, Madison, WI, USA) following the manufacturer’s instructions. All transfection experiments were performed in triplicate. For all *in vitro* cellular experiments (except the MTS assay), the optimized levels of miR-199-3p mimic and mimic NC were 50 nM, while those of the inhibitor and inhibitor NC were 200 nM.

### Prediction of target genes for miR-199-3p and luciferase assay

Genes specifically affected by miR-199-3p targeting were explored by analyzing a combination of *in silico* and transcriptome profiling data. Genes regulated by miR-199-3p were listed using TargetScan (release 7.2) and the miRTarBase database (release 7.0). Genes were obtained from our previous transcriptome sequencing data which are available in the public database (https://www.ncbi.nlm.nih.gov/bioproject/PRJNA506334). The screened target genes of miR-199-3p were categorized via KEGG enrichment analysis in KOBAS 3.0 to further narrow the scope. The wild-type (Wt) or mutant-type (Mut) miR-199-3p target sites in goose *ITGB8* 3′ UTR were obtained by PCR amplification (95°C 5 min, 85°C 5 min, 75°C 5 min, 70°C 5 min). Then, the Wt or Mut sequences were inserted into the pmirGLO Vector (Promega). The vectors were transfected into the goose primary GCs in the presence or absence of mimic or mimic NC using ViaFect^™^ reagent. Cells were lysed at 36 h after transfection and the luciferase activity was analyzed using the Dual-Luciferase Reporter Assay Kit (Promega).

### Cell proliferation and cell migration assays

GCs were transfected with 30 nM, 50 nM and 100 nM mimics, as well as 50 nM, 100 nM, and 200 nM inhibitor, respectively. Cells were plated in 96-well plates at 3×10^5^ cells/well. After 48 h, cell proliferation was determined by MTS assay according to the manufacturer’s instructions, using the CellTiter 96 Aqueous One Solution Cell Proliferation Reagent (Promega). All experiments were performed in replicates of 10. Cell migration was determined by the wound healing assay. Here, goose primary GCs were seeded into 6-well plates at 3×10^5^ cells/well and cultured until confluent. A 10 µl pipette tip was used to make a straight scratch in the middle of each 6-well plate. Then, 50 nM miR-199-3p mimic or 200 nM miR-199-3p inhibitor and corresponding concentrations of NC were transfected into the GCs. After continuous incubation for 18 h, 24 h, and 40 h, the gap widths of scratch re-population were recorded under the microscope at 10 X magnification (Leica DMi8, Wetzlar, Germany) by measuring ten consistently sized areas per well. All the procedures were performed in triplicate, and the data of three independent experiments were analyzed.

### The determination of GC-secreted hormone levels

Progesterone (P4), Anti-Mullerian hormone (AMH) and estradiol (E2) are excreted by GCs. Thus, goose P4, AMH and E2 levels in the supernatant of the cultured GCs were examined using a commercially available ELISA kit, performed according to the manufacturer’s instructions (HengYan Bio., Shanghai, China). Briefly, microtiter plate wells were coated with purified goose antibodies specific for either P4, AMH or E2. The cell culture supernatant was added to the wells followed by a corresponding HRP-labeled antibody to form an antibody-antigen-enzyme-antibody complex. TMB substrate solution was subsequently added and the HRP enzyme-catalyzed, colorimetric reaction was terminated by the addition of a sulfuric acid solution. The color change was measured spectrophotometrically at a wavelength of 450 nm. The concentration of goose P4, AMH and E2 in the samples was finally determined by comparing the optical density value of the samples to the standard curve.

### qRT-PCR analysis of *in vivo* and *in vitro* GCs

Total RNA was extracted from the collected *in vivo* goose and *in vitro* transfection samples using Trizol reagent following the manufacturer’s protocol (Invitrogen, Carlsbad, CA, USA). The RNA concentration and integrity were determined using a NanoDrop 2000 spectrophotometer (Thermo Scientific, Wilmington, DE, USA). To detect and quantify the expression level of miR-199-3p in the granulosa layers, we utilized the Hairpin-it^™^ microRNA qRT-PCR Quantitation Kit (GenePharma, Shanghai, China) following the manufacturer’s protocol. Briefly, the thermal cycling conditions were as follows: 25°C for 30 min, 42°C for 30 min, 85°C for 5 min (reverse transcription reaction); pre-denaturation at 95°C for 3 min, followed by 40 cycles of 95°C for 12 s and 62°C for 40 s (PCR reaction). In addition, mRNA expression levels of *ITGB8, MMP9, MMP15, COL4A1, LAMA1, PCNA, Bcl-2, CASP3, TP53, CCND1, CCND2, SREBP, STAR, 3βHSD* and *FSHR* in the supernatant of cultured GCs were quantified using GoTaq® qPCR Master Mix Kit (Promega) as previously described [29]. All miRNA and mRNA data were normalized by using U6 and *GAPDH* as housekeeping genes, respectively. Relative expression values were calculated using the 2^-ΔΔCt^ method (Livak and Schmittgen, 2001). The primer sequences can be found in Supplementary Tables S2.

### Statistical analysis

Statistical analysis was performed using IBM SPSS Statistics v. 19.0 (IBM Corp., Armonk, NY, USA). All values were presented as the mean ± SD and a *P*-value below 0.05 was considered statistically significant. The area of cell migration was calculated by Image-Pro Plus software.

## RESULTS

### Involvement of miR-199-3p and ECM-related genes during follicle selection

Our published miRNA sequencing data obtained from goose ovarian follicle GCs before (4-6 mm and 8-10 mm in diameter) and after selection (F5) [19] revealed that miR-199-3p was differentially expressed throughout the process of follicular selection (Fig.1A, pink solid column). A higher level of expression appeared in 4-6 mm diameter pre-hierarchical follicles, yet a lower level of expression occurred in follicles of 8-10 mm diameter, then a significant rise was found in F5 hierarchical follicles (*P<*0.05). miR-199-3p levels in the granulosa layers at different stages also followed a similar trend as confirmed by qRT-PCR (Fig.1A, white hollow column). The miR-199-3p level gradually decreased in 2-4 to 8-10 mm diameter pre-hierarchical follicles (*P<*0.05), after which a significant increase occurred in F5-F3 hierarchical follicles (*P<*0.05), and it then decreased again in F2 and F1 hierarchical follicles (*P<*0.05). In short, miR-199-3p levels fell in pre-hierarchical follicles from 2-4 to 8-10 mm diameter (before follicle selection) and rose in the F5-F3 hierarchical follicles (after follicle selection). Thus, we postulated that miR-199-3p played an important role in the process of follicular development, especially follicular selection.

**Fig.1.**
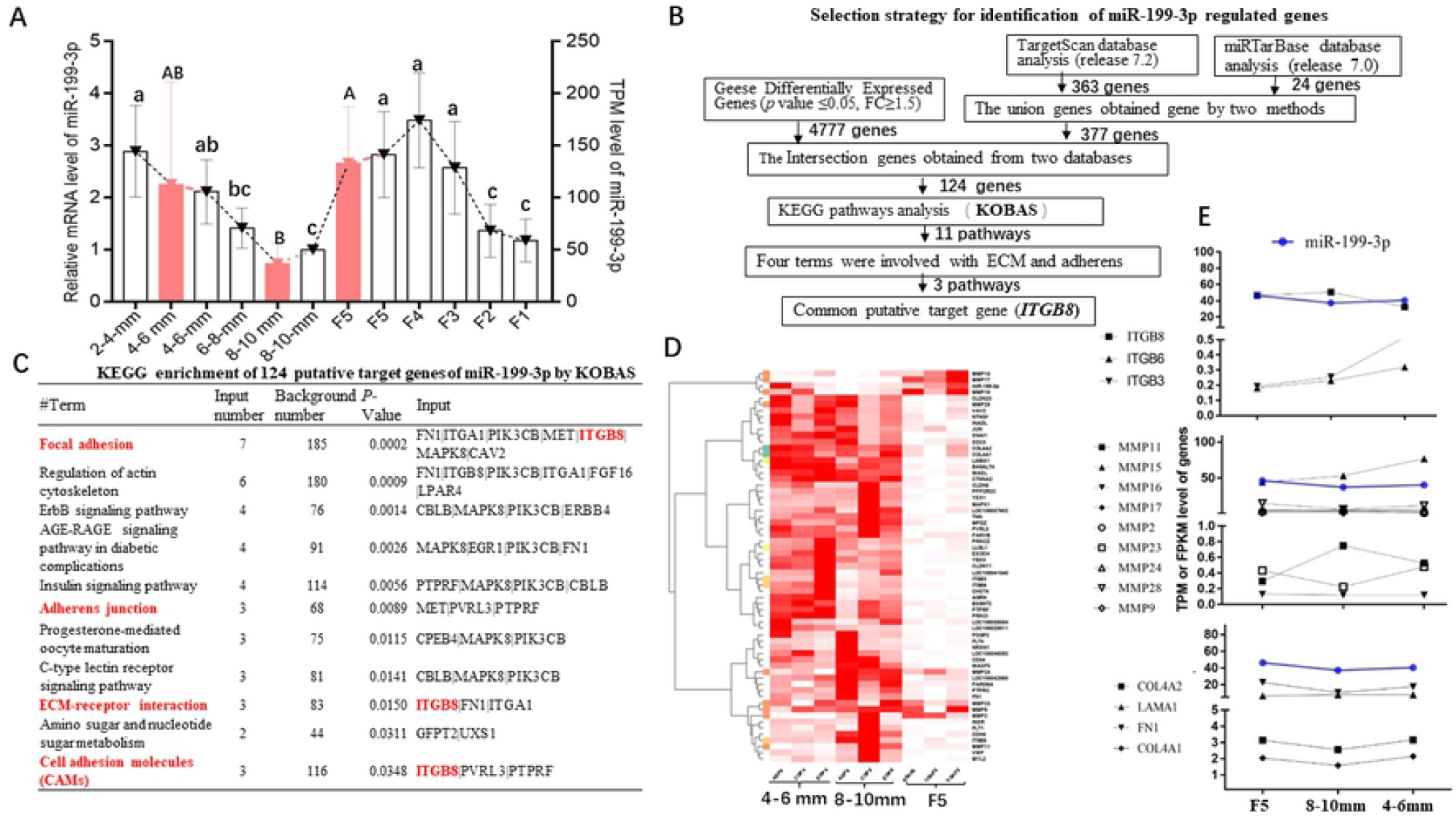
Expression, target gene prediction and the relationship with ECM-related genes of miR-199-3p. (A) The expression of miR-199-3p at different developmental stages in the granulosa layer by sequencing (pink solid column) and qRT-PCR (white hollow column). The letters at the top of each bar represent the significant differences in miR-199-3p gene expression between the various stages (*P*<0.05). TPM levels of miR-199-3p are quoted from our previous study [19]. (B) Flow chart for identification of miR-199-3p target genes. Analysis using TargetScan and the miRTarBase database showed that 363 and 24 genes had putative binding sites for miR-199-3p, respectively; 377 genes were obtained by combining these two methods. Then, 377 genes and 4777 differentially expressed goose genes (NCBI BioProject PRJNA506334) were intersected to obtain 124 genes. Finally, these 124 genes were categorized by GO and KEGG enrichment analysis using the KOBAS (3.0), and 4 of the 11 pathways identified were found to be associated with Focal adhesion, Adherens junctions, ECM-receptor interactions, and cell adhesion molecules (CAMs). (C) KEGG enrichment of 124 putative target genes of miR-199-3p. *ITGB8* was the common gene of 3 pathways (Focal adhesion, ECM-receptor interactions, and CAMs) and may be regulated by miR-199-3p. (D) Heatmap of miR-199-3p and 62 ECM-related genes. (E) The trend chart of miR-199-3p and part ECM-related genes.

To examine the role by which miR-199-3p functions, candidate target genes of miR-199-3p were identified using a combination of TargetScan (Release 7.2) and the miRTarBase (Release 7.0) database and transcriptome data from goose granulosa layers [19]. The specific strategy for selection of target genes is shown in Fig.1B. Our analysis revealed that *ITGB8* was the collective putative target gene of miR-199-3p (Fig.1C). ITGB8 is involved in multiple pathways in relation to cell adhesion (Fig.1C) and is an ECM adhesion receptor [30]. Therefore, utilizing our previous transcriptome sequencing data from the granulosa layer tissues of geese [19], we analyzed the clustering pattern of 62 ECM-related genes including *ITGBs, LAMA1, COL4As* and *MMPs*, as well as the relationship between these genes and miR-199-3p (Fig.1D and Fig.1E). The results showed that these ECM-related genes were mainly clustered into three classes (Fig.1D). The first class contained miR-199-3p and some of the MMPs; the levels of these genes were lower in follicles of 8-10 mm diameter compared to follicles of 4-6 mm in diameter and F5. The second class, including *LAMA1* and *COL4As*, comprised mostly of ECM-related genes; the differences in these genes were more pronounced during follicular selection (*i*.*e*. F5 *vs*. 8-10 mm group) compared to before follicle selection (*i*.*e*. 8-10 mm *vs*. 4-6 mm group). The third class included *ITGB8* and contrary to the first class, showed an opposing trend to miR-199-3p (Fig.1D and Fig.1E). Therefore, according to the changes of these ECM-related genes and miR-199-3p, we postulated that the putative miR-199-3p target gene *ITGB8* and other ECM-related genes play important roles in regulation of avian follicular selection.

### miR-199-3p suppressed goose granulosa cell viability and migration

Previous evidence has shown that miR-199 exerts a suppressive effect on cell viability and migration in several type of cancer cell [22], although the effect of miR-199 on ovarian follicles has not been reported. We performed gain-of-function and loss-of-function studies using goose GCs transfected with either a mimic or inhibitor of miR-199-3p, respectively. MTS assays revealed a significant inhibition of cell viability when transfected with 30 nM and 50 nM of mimic (*P*<0.05 or *P<*0.01; Fig.2A) compared with their mimic-NCs, yet a significant promotion of cell viability when transfected with 200 nM inhibitor versus the corresponding NC (*P<*0.01; Fig.2B). Moreover, wound-healing assays demonstrated that miR-199-3p overexpression could suppress cell migration activity (*P<*0.05; Fig.2C) yet loss of miR-199-3p could promote migration after 18h or 24h of transfection compared to their corresponding controls (*P<*0.05; Fig.2D). Taken together, these data demonstrated that miR-199-3p suppressed *in vitro* viability and migration of goose GCs.

**Fig. 2.**
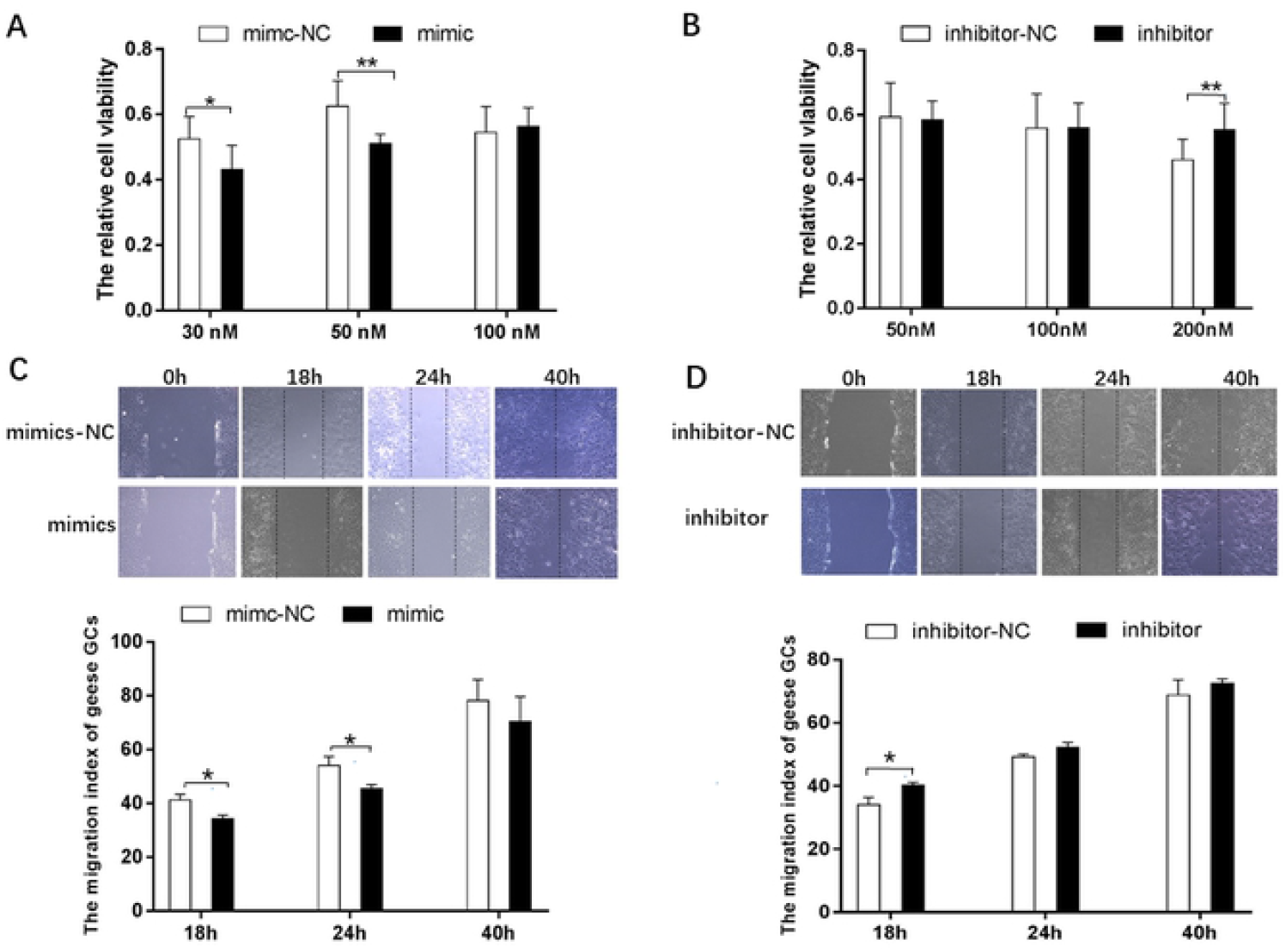
miR-199-3p mimics inhibited cell viability and migration of goose GCs. (A-B) Effect of miR-199-3p on viability of goose GCs. (C-D) Images and histogram acquired at 0, 18, 24, and 40 h in the wound-healing assay. Goose GCs shown were transfected with miR-199-3p mimics, mimics-NC (C) and inhibitor, inhibitor-NC (D). The dotted lines define the areas lacking cells. The area of cell migration was calculated by Image-Pro Plus software. The data are expressed as the mean ± SD; n = 5. \**P* < 0.05 and \*\**P*<0.01 versus the NC.

### miR-199-3p promoted steroidogenesis and the expression of key genes for steroidogenesis

Due to the important effects of steroidogenesis on GCs, we further demonstrated the role of miR-199-3p in goose GC steroid production. We found that miR-199-3p significantly increased P4 (*P*<0.05) but not E2 and AMH concentrations (Fig. 3A), and significantly elevated the levels of *STAR* and *3βHSD* but not *SREBP* and *FSHR* (*P*<0.05; Fig. 3B) in the supernatant of cultured GCs when compared to the control. These data suggest that miR-199-3p participates in goose GC steroidogenesis.

**Fig.3.**
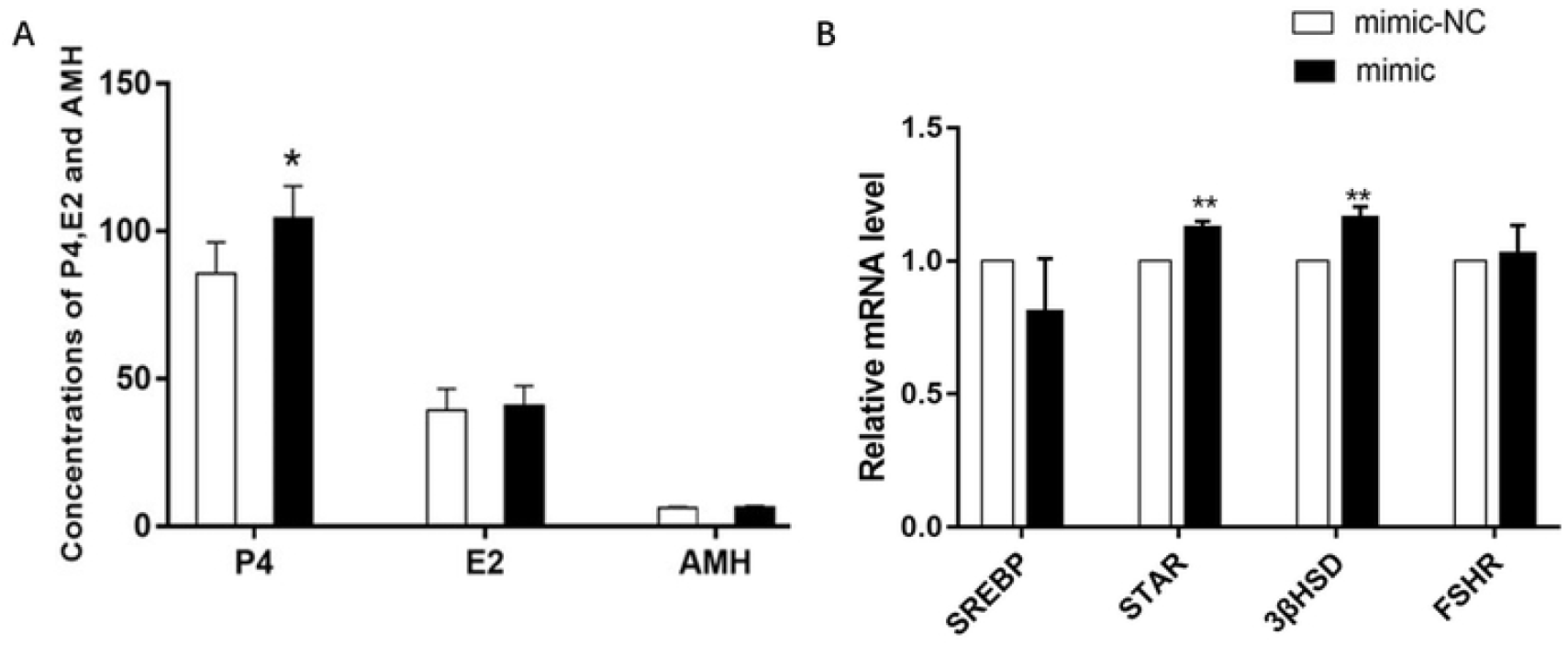
The expression levels of hormones (A) and steroidogenesis-related genes (B) after miR-199-3p overexpression. Data are presented as the mean ± SD of three independent experiments. *GAPDH* was used as the reference gene for qRT-PCR data normalization. * *P*<0.05. \*\**P* < 0.01. P4, E2 and AMH are abbreviations for progesterone, estradiol, and anti-Mullerian hormone, respectively.

### MiR-199-3p directly targeted *ITGB8* and can regulate expression of ECM-related genes

We predicted that miR-199-3p binds to position 264-270 of the *ITGB8* 3′ UTR based on the existence of a highly conserved site among different species (Fig.4A-4C). We found that luciferase activity was significantly reduced in 293T cells by co-transfection with miR-199-3p mimics and the vectors carrying the *ITGB8* 3′ UTR -WT reporter but not those carrying the MUT (Fig.4D). These data indicate that miR-199-3p directly binds to sites in the 3′ UTR of *ITGB8* mRNA. Furthermore, we co-transfected GCs from goose pre-hierarchical SWF follicles with miR-199-3p mimics and *ITGB8* 3′ UTR-WT reporter, and the effects of miR-199-3p overexpression on the levels of ECM-related genes were examined by qRT-PCR. The result showed that miR-199-3p overexpression significantly downregulated the expression levels of *ITGB8, MMP9* and *MMP15* (*P*<0.01) yet upregulated *COL4A1* and *LAMA1* expression (*P*<0.05; 4E). In addition, the levels of proliferation and apoptosis genes, including *PCNA, Bcl-2, Caspase 3, CCND1*, and *CCND2* (except for *TP53*) had not changed (Fig. 4F). In combination with the foregoing effects of miR-199-3p on goose GCs, the obtained results indicate that miR-199-3p may be involved in cell viability, migration, or steroidogenesis by directly targeting *ITGB8* or indirectly regulating other ECM-related genes during avian follicular selection.

**Fig. 4.**
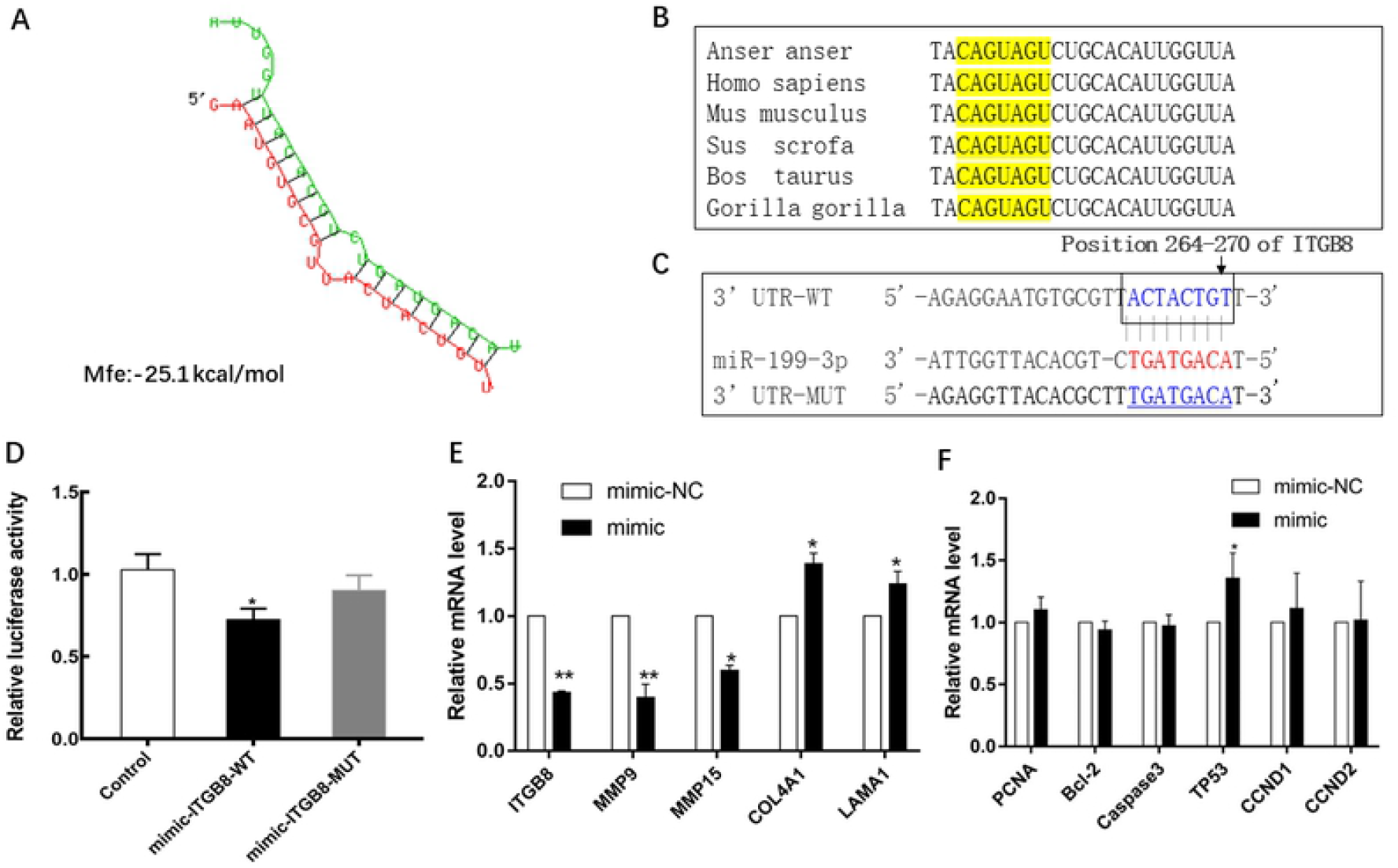
*ITGB8* is a direct target of miR-199-3p. (A) The schematic of miR-199-3p depicting binding to its targeted gene using RNAhybrid. The green sequence is the miR-199-3p mature sequence; the red sequence is the miR-199-3p target gene. (B) The miR-199-3p sequence is shown to be highly conserved among species (yellow background area). (C) miR-199-3p binding sites in the 3′ UTR of *ITGB8*. Dual luciferase reporter assays using vectors encoding putative miR-199-3p target sites (red) of the *ITGB8* 3′ UTR (positions 264-270) for wild type or mutant type (blue). (D) Luciferase assay of 293T cells co-transfected with firefly luciferase constructs containing the *ITGB8* 3′ UTR-WT, MUT and miR-199-3p mimics, mimics-NC, miR-199-3p inhibitor or inhibitor NC, as indicated (n=5). (E) The expression levels of genes associated with ECM after miR-199-3p overexpression in goose GCs. (F) The expression levels of proliferation and apoptosis genes after miR-199-3p overexpression in goose GCs. Normalized data were calculated as ratios of Renilla/firefly luciferase activities. Data are presented as the mean ± SD of three independent experiments. * *P* < 0.05, ** *P* < 0.01.

## DISCUSSION

The present study is the first to investigate the function of miR-199-3p on GCs in birds. Firstly, a significant change in miR-199-3p in the granulosa layer of geese was observed before and after follicular selection in our previous sequencing data and verified by qRT-PCR, suggesting that miR-199-3p plays an important role in the regulation of follicular selection. Further, we identified *ITGB8* as the putative target gene of miR-199-3p and predicted that ECM-related genes play an important role during the process of follicular selection by analyzing *in silico* database and geese transcriptome data. To clarify the molecular regulatory mechanism of miR-199-3p, we confirmed ECM-related gene *ITGB8* as a direct target of miR-199-3p by dual-luciferase reporter assay. Additionally, the overexpression of miR-199-3p in the cultured GCs of geese pre-hierarchical follicles suppressed two ECM-related genes (*MMP9* and *MMP15)* yet enhanced the expression of another two ECM-related genes (*COL4A1* and *LAMA1)*. Previous research has shown that MMPs, collagen type IV, and fibronectin are ECM constituents and play important and dynamic roles in cell migration, survival, and steroidogenesis [11, 31]. MMPs are the enzymes that break down ECM substrates such as collagen type IV and laminin to facilitate tissue remodeling [13]. Degradation occurs to unpack the tissue and ECM of the follicle to release the oocyte [9] and to facilitate deposition of large amounts of yolk precursors via an extensive vasculature during avian follicular selection [32, 33]. MMPs play key roles in the follicular ECM transformation associated with follicle development and maturation [10, 34, 35]. Therefore, in our present study, the rise of *COL4A1* and *LAMA1* and the fall of *MMP9* and *MMP15* implies that miR-199-3p overexpression reduced the degradation effect of MMPs on *COL4A1* and *LAMA1* genes during goose follicular selection to decrease follicular ECM remodeling. Together, the present findings showed that miR-199-3p may be involved with follicular ECM remodeling by targeting *ITGB8* and regulating ECM-related genes during avian follicular selection.

On the other hand, transfection of miR-199-3p mimics and inhibitors into goose pre-hierarchical follicle GCs revealed a suppressive effect of miR-199-3p on the viability and migration of GCs, as demonstrated by the MTS and wound-healing assays. To date, the function of miR-199-3p on ovarian follicles has rarely been studied. Some research found that miR-199 suppressed cancer cell proliferation, migration, and invasion [21, 22, 36]. The findings of the present study also suggested that miR-199-3p plays a similar inhibitory role in avian ovarian follicles. Furthermore, it has corroborated the importance of ECM remodeling by MMP-driven pericellular proteolysis in facilitating cell migration [37]. Similarly, cell migration has an important role in supporting the morphological changes required for extensive tissue remodeling to facilitate maturation and release of oocytes [38, 39]. Thus, our results suggest that miR-199-3p is involved in the follicular remodeling mediated by migration of GCs during follicular selection. Meanwhile, the inhibitory effect of miR-199-3p on the migration of goose GCs may additionally help to explain to a certain extent the role of *MMP9* and *MMP15* downregulation after miR-199-3p overexpression.

Furthermore, our data revealed that P4 and the expression levels of key genes required for P4 production (*STAR* and *3βHSD*) were significantly increased after miR-199-3p overexpression, indicating that miR-199-3p could modulate steroidogenesis in goose GCs. To date, no research on the role of miR-199-3p on follicular function and steroidogenesis has been identified. However, it has been suggested that collagens (*COL4A1* and *COL4A4*), laminin, and fibronectin increase cellular attachment and P4 production in cultured porcine GCs [40]. In turn, *MMP1* and *MMP3* mRNA expression has been shown to be stimulated by P4, FSH, and estrogen in chicken GCs [10]. These studies indicate collagen, laminin, and MMPs could affect steroidogenesis. In the present study, miR-199-3p overexpression promoted laminin and collagen type IV mRNA expression and increased P4 secretion and mRNA expression of *STAR* and *3βHSD*, which are required for P4 production. Furthermore, *STAR* and *3βHSD* are markers for GC differentiation and are closely related to steroidogenesis after follicular selection [19]. Thus, our results suggest that miR-199-3p can modulate P4 production during goose follicular selection.

## CONCLUSIONS

The present study demonstrated that miR-199-3p was differentially expressed in goose ovarian follicles before and after follicular selection. Furthermore, miR-199-3p was shown to suppress viability and migration, as well as promote both the secretion of P4 and the expression of key genes for P4 production in cultured GCs from goose follicles prior to selection. Finally, miR-199-3p was found to directly target the ECM-related gene *ITGB8* and regulate other ECM-related genes, including *MMP9, MMP15, COL4A1*, and *LAMA1* in cultured GCs from goose pre-hierarchical follicles. These findings indicate that miR-199-3p may be involved in granulosa cell viability, migration, and steroidogenesis in goose ovarian follicles prior to selection by targeting *ITGB8* and regulating other ECM-related genes.

